# DREIMT: a drug repositioning database and prioritization tool for immunomodulation

**DOI:** 10.1101/2020.06.24.168468

**Authors:** Kevin Troulé, Hugo López-Fernández, Santiago García-Martín, Miguel Reboiro-Jato, Carlos Carretero-Puche, Jordi Martorell-Marugán, Guillermo Martín-Serrano, Pedro Carmona-Sáez, Daniel González-Peña, Fátima Al-Shahrour, Gonzalo Gómez-López

**Affiliations:** Bioinformatics Unit, Spanish National Cancer Research Centre (CNIO), Madrid 28029, Spain; SING Research Group, CINBIO - Biomedical Research Centre (University of Vigo), IIS Galicia Sur - Galicia Sur Health Research Institute (SERGAS-UVIGO); Department of Computer Science, University of Vigo, Ourense, Spain; Centre for Genomics and Oncological Research (GENYO), Granada, Spain

## Abstract

**Motivation:** Drug immunomodulation modifies the response of the immune system and can be therapeutically exploited in pathologies such as cancer and autoimmune diseases.

**Results:** DREIMT is a new hypothesis-generation web tool which performs drug prioritization analysis for immunomodulation. DREIMT provides significant immunomodulatory drugs targeting up to 70 immune cells subtypes through a curated database that integrates 4,960 drug profiles and ~2,6K immune gene expression signatures. The tool also suggests potential immunomodulatory drugs targeting user-supplied gene expression signatures. Final output includes drug-signature association scores, FDRs and downloadable plots and results tables.

**Availability:** http://www.dreimt.org

## 1 Introduction

Immune system dysregulations have been related to a wide spectrum of complex diseases such as autoimmune disorders and cancer. Autoimmunity processes are clinically diverse and they are distinguished by an immune-mediated attack on the body’s own tissues led by self-reactive B and T cells (Rose *et al*. 2016). In cancer, the intercellular signalling between the malignant cells and certain immune cell subpopulations (e.g. regulatory T cells (Tregs) and tumor-associated macrophages (TAMs)) contributes to tumor microenvironment immunosuppression fostering cell proliferation and tumor evasion (Stockis *et al*. 2019; Linde *et al*. 2018). The presence of Tregs and TAMs subpopulations in tumor microenvironment has been also correlated with cancer poor prognosis in contrast to better prognosis shown by those tumors with high rate of tumor-infiltrated CD8+ T cells (Fridman *et al*. 2017). This has led to propose immune cells targeting as a therapeutic strategy. For instance, some studies have shown that activity of specific immune cell populations can be targeted using systemic or non-systemic approaches (e.g. drug-conjugated nanoparticles) thus enhancing the outcome of some tumors and autoimmune diseases (Lu *et al*. 2020; Riley *et al*., 2019; Genovese *et al*., 2016). Recently, drug-mediated immunomodulation has been proposed as a therapeutic approach in patients with COVID-19 associated cytokine storm (Richardson P *et al*., 2020). This highlights the interest in developing new methodologies to propose immunomodulatory therapeutic strategies capable of selectively affect the function of specific immune cells. Here we introduce DREIMT (Drug REpositioning for IMmune Transcriptome), a tool for hypothesis generation of drugs capable of modulating (boosting or inhibiting) the immune cells activity.

## 2 Methods and features

DREIMT integrates ~2.6k immune gene expression signatures with a collection of 4690 consensus drug profiles (Perales-Patón *et al*., 2019) from the catalog of drug perturbations expression profiles in cancer cell lines from the LINCS L1000 dataset (Subramanian *et al*., 2017) (Figure 1). Immune signatures were obtained from published studies and public databases covering up to 70 immune cell subtypes (Suppl. Materials S1-3, T1). DREIMT performs a fgsea to test the significance of the immune signature enrichment across the ranked genes of each drug profile (Korotkevich *et al*., 2020). The fgsea enrichment scores are used to calculate the Drug Prioritization score (⊤) as described elsewhere (Subramanian et al., 2017), comparing the obtained enrichment score for a particular signature to the rest of immune signatures. The ⊤ score allows the prioritization of drugs to target immune cells. DREIMT also includes a Drug Specificity Score (DSS) that summarises the cell-specificity of a given drug across multiple cancer cell lines (Hodos *et al*., 2018; Suppl. Materials S4-6). Significant drug-immune signature associations (|⊤|>80) are stored in the DREIMT database (DREIMTdb) accessible through a RESTful API. We have validated >20 DREIMT hypothesis by scientific literature (Suppl. Materials S7, T2).

**Fig. 1.**
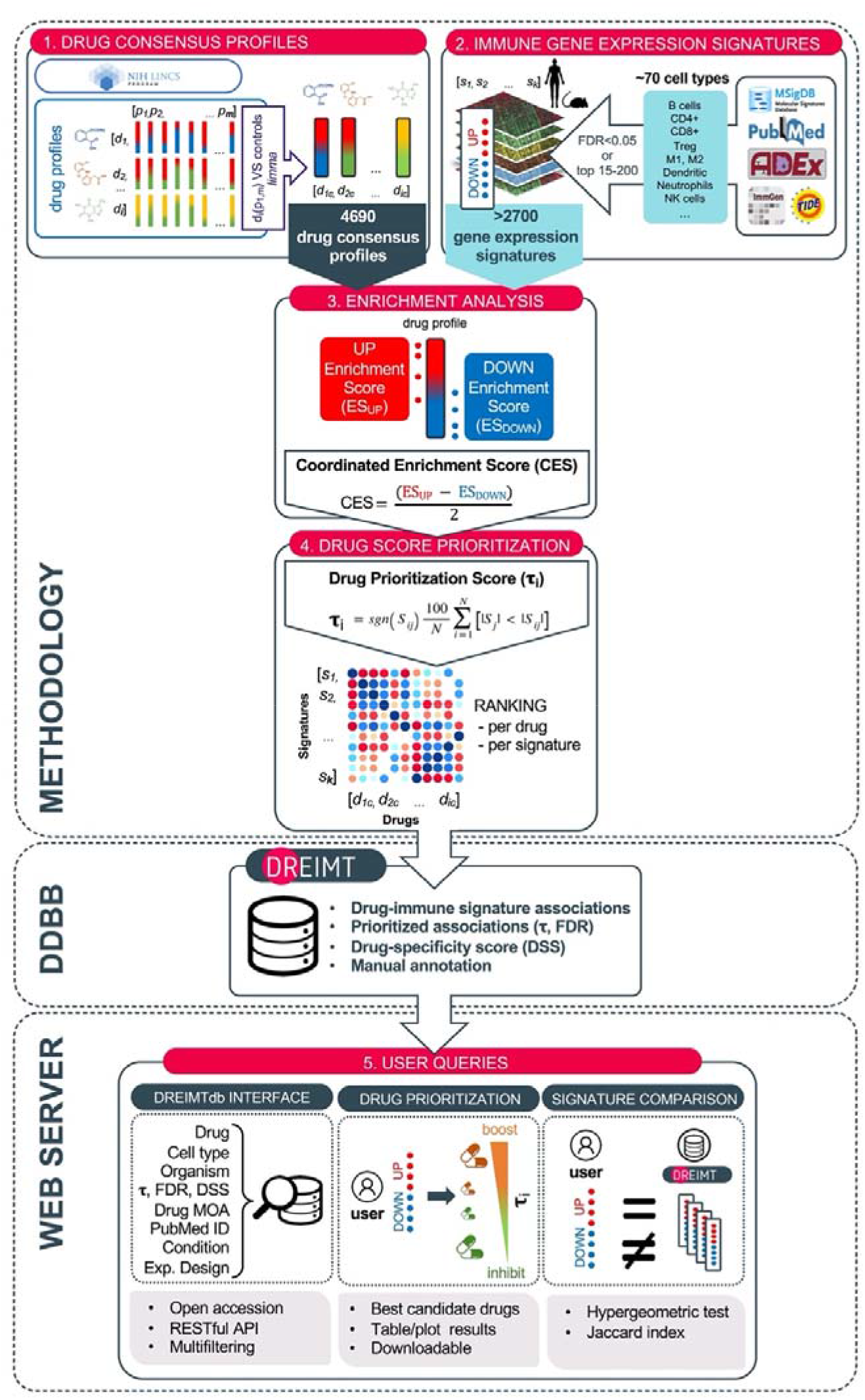
DREIMT workflow.

## 3 DREIMT web server

DREIMT web server functionalities include DREIMTdb interface, drug prioritization and signature comparison tools (Figure 1). DREIMTdb interface allows an easy interrogation of the drug-immune signature associations allowing multi-filtering for advanced queries. Drug prioritization functionality allows the search for the best candidate drugs to boost or inhibit the user-defined immune gene expression signature of interest. Drug results are prioritized by ⊤ and DSS and filtered by statistical significance (FDR<0.05). Final outputs are displayed as fully downloadable tables and plots. Additionally, users can compare their own gene signatures against full DREIMTdb contents. As a result, the tool provides the similarity score (jaccard index) and FDR value per each signature compared (Suppl. Materials S8,9). Implementation details and links to the source code repositories, are provided in the Suppl. Materials S10.

## 4 Conclusions

Drugs affect the biology and activity of the immune system, however such interactions are often poorly understood. We have developed DREIMT, a tool to allow users to generate hypotheses and explore novel druggable targets across the immune system, thus supporting drug repositioning leveraging transcriptomics data. DREIMT is fully accessible at http://www.dreimt.org.

## Acknowledgements

José Antonio López for the fruitful discussions for the development of the DREIMT concept. BU staff for beta testing and DREIMTdb curation.

