## Supplementary Material for "DREIMT: a drug repositioning database and prioritization tool for immunomodulation"

### INDEX

**Page**

**Index**……………………………………………………………………………… 2
**S1.** Immune signatures collection………………………………………………… 3
**S2.** Immune signatures annotation……………………………………………….. 3
**S3.** Drug-associated gene expression profiles………………………………….. 4
**S4.** Estimation of drug profile-immune gene expression associations……….. 4
**S5.** Drug prioritization score…………………………………………………… 5
**S6.** Drug annotation………………………………………………………………... 6
**S7.** DREIMT database description……………………………………………... 7
**S8.** DREIMT drug prioritization tool……………………………………………. 7
**S9.** DREIMT signature comparison tool………………………………………… 8
**S10.** DREIMT software implementation details and availability……………… 8
**T1.** Cell types and subtypes available in DREIMTdb………………………….. 10
**T2.** Examples of DREIMT hypotheses confirmed in scientific literature…….. 12
**References**……………………………………………………………………... 13

* * * * * * * * * *

#### S1. Immune signatures collection.

Immune gene expression signatures (2,687) were obtained from the following sources: MsigDB C7 collection (2,436) (Godec et al., 2016), ImmGen (173) (Heng et al., 2008), ADEx (35 signatures) (https://adex.genyo.es) and from publicly available gene expression datasets (43 signatures).

Gene expression signatures for ImmGen, ADEx and public datasets were obtained using *limma* Bioconductor's package (Ritchie et al., 2015). These signatures of upregulated and downregulated differentially expressed genes (DEG), were selected under the criteria of significance (FDR < 0.05) and ranked by log2FC with a minimum of 15 genes and a maximum of 200 genes. Genes were annotated following the HUGO Gene Nomenclature Committee format (Yates et al., 2017).

#### S2. Immune signatures annotation.

The DREIMT immune signature collection comprises 1,667 *Mus musculus* signatures and 1,020 corresponding to *Homo sapiens*. The collection was manually annotated to ease DREIMT results interpretation. Immune cells were annotated at two levels: i) cell type and ii) cell subtype. The "cell type" level includes 17 major immune cell types while "cell subtype" level second comprises 70 immune cell subtypes see Supplementary Table 1 for details. Additionally, the immune signatures were manually annotated with: experimental conditions undergone by the cells (type of stimulation, gene knock-outs and disease models), experimental design (patient, *in vivo*, *in vitro*), organism (*Homo sapiens, Mus musculus*) and PubMed ID source of the original experiment and the immune comparison.

The 2,687 immune signatures integrated in DREIMT represent a wide spectrum of conditions due to the large variation in experimental and biological designs. Changes in lineage, immune cell activation or developmental status and perturbation effects are represented.

**S3. Drug-associated gene expression profiles.**

Drug profiles were obtained from the The Library of Network-Based Cellular Signatures (LINCS) L1000 data set (Broad Institute LINCS Center for Transcriptomics (1U54HL127366)) from the Connectivity Map (CMap) project. LINCS provides a large-scale catalogue of transcriptional responses to pharmacological and genetic perturbations upon a large panel of cell lines (Subramanian et al., 2017).

Consensus drug profiles were generated from LINCS L1000 level 3 data (quantile $\log_2$FC normalized gene expression profiles) using limma (v 3.24.15, R package) by comparing the differential gene expression between treated and untreated control experiments for a given perturbation (drug) using additive linear model to avoid batch effects and individual cell lines drug responses. These consensus drug profiles represent the consensus cellular-specific perturbation for 12,434 genes ranked by expression fold change ($\log_2$FC) and they are expected to represent the common transcriptional changes induced across different cell lines, replicates and drug concentrations (Perales-Patón et al., 2019). A total of 4,690 consensus drug profiles were included in DREIMT, of which 3,583 correspond to unique drug compounds.

**S4. Estimation of drug profile-immune gene expression associations.**

Statistical relevance of the drug profile-immune expression signature associations were calculated using the Kolmogorov-Smirnov test implementation available in Bioconductor's fast gene set enrichment analysis package (FGSEA; version 1.0.0) (Korotkevich et al., 2019). We employed 1000 random permutations for error estimation.

Two different procedures are followed to calculate the Enrichment Score (ES):

a) FGSEA ES calculation for a gene set defined based on *a priori* knowledge (e.g. genes functionally related to the immune system activity) or a gene list (e.g. statistically significant genes obtained from immune multi-omics experiments).

b) Pairwise ES calculation for immune gene expression signatures. Here FGSEA ES is calculated for upregulated ($ES_{up}$) and downregulated ($ES_{down}$) genes separately. The $ES_{up}$ and $ES_{down}$ obtained are then used to generate a coordinated enrichment score (CES). The CES is created by subtracting the $ES_{down}$ from the $ES_{up}$ and dividing this by two (i.e. CES= $[ES_{up} - ES_{down}]/2$). This CES ranges from 1 to -1 representing the enrichment coordination for the upregulated and downregulated genes across a given drug profile.

Both ES and CES represent the association between a given consensus drug profile and an immune geneset/genelist or immune expression signature respectively.

P-value estimation is calculated using an adaptive multi-level split Monte-Carlo scheme (see *fgsea* package documentation for details). To account for multiple hypothesis testing, the estimated p-values are adjusted using Benjamini & Hochberg False Discovery Rate (FDR) correction (Benjamini Y. and Hochberg Y., 1995). Those drug profile-immune signature associations showing FDR <0.05 were considered statistically significant.

**S5. Drug prioritization score.**

Drug prioritization score (τ, tau) is calculated following the approach adopted by the LINCS L1000 team (Subramanian et al., 2017).

This score ranging from -100 to 100 measures: i) How similar is a particular drug-immune gene expression association compared to the rest of associations for a given drug and ii) how similar is a particular drug-immune gene expression association compared to all the associations calculated for a given immune signature.

More specifically, an ES (or CES) matrix for all drug profile-immune signature associations (i.e. immune signatures in rows and drug profiles in columns) is firstly generated. Drug prioritization score calculation is performed by normalizing a given ES (or CES) by the mean values of the drug (column) and by the mean values of the signature (row) to which that particular drug-immune gene expression association belongs (only values with the same association score sign are used). Normalized association scores are column scaled (ranging from -1 to 1) and then column standardized ranging from -100 to 100.

DREIMT results are prioritized by tau value. A tau score >|80| is considered as strong hypotheses for the drug profile-immune signature associations.

**S6. Drug annotation.**

Drugs status and mode of action (MOA) was retrieved from the Drug repurposing hub (version 20200324). For some drugs with no information available, drug status, and MOA were manually annotated using the literature and public resources. In summary, 64% of drugs have MOA annotated, 53% a gene target, 43% a DSS and all drugs have been included with drug status information (approved, experimental or withdrawn).

Expression response to a drug exposure can be highly cell-specific (due to drug target transcriptional changes) or induce very similar expression across cell types. To consider this issue, DREIMT includes a Drug Specificity Score (DSS) adapted from Hodos et al. (Hodos et al. 2018) that summarises the cell-specificity of a given drug across multiple cancer cell lines. DSS has been inverted and scaled using the originally reported values to ease the interpretation, for this purpose the DSS in DREIMT range from 0 to 1. High DSS value indicates that the transcriptomic effect of the drug in LINCS is found to be similar across all cancer cell lines in LINCS database, while DSS closer to 0 indicates that the drug effect is cancer cell line specific. We assume that since some drugs induce similar gene expression signatures across cancer cell lines (high DSS value) their effects can be more easily extrapolated to immune cells than those produced by drugs with very specific effects described in cancer cell lines (low DSS). DSS is available in DREIMT for 2,044 of drug profiles.

#### S7. DREIMT database description.

DREIMT database (DREIMTdb) contains all the drug-immune signature associations showing |tau score| > 80. DREIMTdb queries can be customized by multi filtering options including drugs, cell types, cell subtypes, experimental conditions, organism, source of the immune signature, interaction type, experimental design, drug status, drug mode of action, etc. DREIMTdb annotations have been manually curated.

DREIMTdb can be queried using DREIMT web tool, but also, through its application programming interface (REST-API) available at http://www.dreimt.org/api. In addition, full DREIMTdb is available as an .RDS file at https://gitlab.com/bu_cnio/DREIMT.

#### S8. DREIMT drug prioritization tool.

DREIMT allows drug prioritization analysis using users-provided immune signatures and genesets. The approach employed to prioritize drugs to target users' immune signatures is described in the sections S4 and S5 of this document.

To launch a drug prioritization analysis in DREIMT users simply have to copy and paste their genes of interest into the boxes provided by the web tool. Alternatively, users can load their gene lists as .CSV and .TSV input files (one gene list per column). Upregulated and downregulated genes from the same experiment can be tested simultaneously, in this case, the columns containing the gene lists must be contiguous (see DREIMT Help for further details). Accepted gene sets size goes from min. 15 to max. 200 genes.

DREIMT drug prioritization results output includes tau score and FDR value for each drug suggested. DSS, MOA and drug status are also included. Drug prioritization full results table can be downloaded as a .CSV file.

**S9. DREIMT signature comparison tool.**

DREIMT signature comparison tool allows comparisons between user-provided gene lists and immune signatures stored in DREIMTdb. Users can perform gene list comparisons against the whole DREIMTdb. Alternatively multi-filtering options allow comparisons against a specific set of DREIMTdb contents.

The overlap between gene lists is tested using Fisher's exact test implementation available in the Bioconductor's GeneOverlap library (R package version 1.23.0). P-values are adjusted using Benjamini & Hochberg False Discovery Rate (FDR) correction. Jaccard index, defined as the number of intersections over the number of unions, is also provided. Full results table can be downloaded as a .CSV file.

**S10. DREIMT software implementation details and availability.**

The DREIMT front-end is implemented using the Angular v9 development framework, the Angular Material v9 library and the Material Dashboard Angular 5 template.

The DREIMT back-end is implemented using Java EE 7 and performs the different data analyses (drug prioritization and similar signatures) using custom R scripts through a docker image. Both the R scripts and the docker image are publicly available in the links provided in the table below.

The communication between front-end and back-end is done using AJAX and JSON. The back-end runs in a WildFly v10.1.0 application server and uses a MySQL database to store the data.

| Software | URL |
| --- | --- |
| Website | http://www.dreimt.org |
| DREIMTdb REST API | http://www.dreimt.org/api |
| DREIMT docker | https://hub.docker.com/r/singgroup/r-dreimt-scripts |
| Data analysis R Scripts source code | https://gitlab.com/bu_cnio/DREIMT |
| Front-end source code | https://github.com/sing-group/dreimt-frontend |
| Back-end source code | https://github.com/sing-group/dreimt-backend |

* * * * * * * * * *

#### T1. Cell types and subtypes available in DREIMTdb.

| Type | Subtype |
|---|---|
| B cell | - |
| B cell | B cell memory |
| B cell | B cell naïve |
| B cell | Plasma cell |
| Basophil | - |
| Dendritic cell | - |
| Eosinophil | - |
| Erythroblast | - |
| Granulocyte | - |
| Hematopoietic stem and progenitor cells | - |
| Hematopoietic stem and progenitor cells | Thymocyte |
| Macrophage | - |
| Macrophage | Macrophage M1 |
| Macrophage | Macrophage M2 |
| Macrophage | Macrophage naïve |
| Mast cell | - |
| MDSC | - |
| MDSC | Monocyte\|MDSC |
| Megakaryocyte | - |
| Monocyte | - |
| Neutrophil | - |
| NK cell | - |
| NK cell | NK naïve |
| Other | - |
| Other | CD80+ cell |
| Other | Cell line |
| Other | Centroblast |
| Other | Centrocyte |
| Other | Consensus signature |
| Other | Endothelial |
| Other | Epithelial |
| Other | Fibroblast |
| Other | Liver |
| Other | Lung |

| | |
|---|---|
| Other | Melanoma |
| Other | Microglial |
| Other | Multiple |
| Other | Nuocyte |
| Other | Plasmablast |
| Other | Plasmacytoid |
| Other | Skin |
| Other | Spleen |
| Other | Splenocyte |
| Other | T2M cell |
| Other | Thymic |
| Other | Thymic stroma |
| PBMC | - |
| PBMC | Myeloid |
| T cell | - |
| T cell | T CD4+ |
| T cell | T CD4+ effector |
| T cell | T CD4+ memory |
| T cell | T CD4+ naïve |
| T cell | T CD8- |
| T cell | T CD8+ |
| T cell | T CD8+ effector |
| T cell | T CD8+ memory |
| T cell | T CD8+ naïve |
| T cell | T cell gamma/delta |
| T cell | T follicular helper |
| T cell | T helper |
| T cell | T helper 0 |
| T cell | T helper 1 |
| T cell | T helper 1 naïve |
| T cell | T helper 17 |
| T cell | T helper 17 naïve |
| T cell | T helper 2 |
| T cell | T naïve |
| T cell | T regulatory |
| T cell | Thymus |

#### T2. Examples of DREIMT hypotheses confirmed by scientific literature.

| DREIMT summary | Signature | UpFDR / DownFDR / Tau | Evidence in scientific literature |
|---|---|---|---|
| **oxaprozin** boosts resting T CD4+ compared to activated T CD4+ effector | GSE13738_RESTING_ VS_TCR_ACTIVATED_CD4_TCELL | 0.032 / 0.010 / 92.28 | Birmingham et al., *Practical Manag. of Pain* 2014; Paccani et al., *JBC* 2002. |
| **antimycin-a** boosts Macrophage compared to Macrophage stimulated with CSF1 | GSE11864_UNTREATED_VS_CSF1_IN_MAC | 0.397 / 0.036 / 99.92 | Jones et al., *Organogenesis*. 2013; Van den Bossche et al., J. *Vis. Exp.* 2015. |
| **meclozine** inhibits Macrophage compared to Macrophage stimulated with CSF1 | GSE11864_UNTREATED_VS_CSF1_IN_MAC | 0.037 / 0.595 / -90.07 | Guo et al., *Front. Pharmacol.* 2017; Huang et al., *Bone Res.* 2017. |
| **mirtazapine** inhibits Macrophage M1 compared to Macrophage M2 | GSE5099_CLASSICAL_M1_ VS_ALTERNATIVE_M2_MACROPHAGE | 0.132 / 0.071 / -90.04 | Almishri et al., *Front. Immunol.* 2019. |
| **sulpiride** boosts Monocyte compared to Macrophage M1 | GSE5099_MONOCYTE_ VS_CLASSICAL_M1_MACROPHAGE | 0.447 / 0.012 / 91.25 | Haskó et al., *Immun. Lett.* 1996. |
| **clonidine** boosts Monocyte vs Macrophage M1 | GSE5099_MONOCYTE_ VS_CLASSICAL_M1_MACROPHAGE | 0.565 / 0.015 / 92.77 | Romero-Sandoval et al., *Brain Behav. Immun.* 2008. |
| **PNU-120596** boosts T regulatory vs T cell | GSE7852_TREG_VS_TCONV | 0.188 / 0.343 / 99.92 | Zhao et al., *J Neuroinflammation* 2017. |
| **ozagrel** boosts T regulatory vs T cell | GSE7852_TREG_VS_TCONV | 0.823 / 0.122 / 93.11 | Cong et al., *Sci. Rep.* 2016; Lim et al., *Int. Immunopharmacol.* 2015. |
| **flunisolide** boosts Macrophage vs Macrophage stimulated with IFNg | GSE1925_CTRL_ VS_24H_IFNG_STIM_MACROPHAGE | 0.05 / 0.31 / 94.93 | Bewing et al., *Eur. J. Clin. Pharmacol.* 1993. |
| **BIX-01294** inhibits T CD8+ stimulated with LCMV and Overexpression (TOX) vs T CD8+ stimulated with LCMV | Tcell_exhaustion_TOX_overexpression_ VS_TOX_control_LCMV | 0.014 / 0.043 / -91.08 | Barili et al., *Nat. Commun.* 2020. |
| **HNHA** boosts Dendritic cell vs Dendritic cell stimulated with LPS | GSE36009_UNSTIM_VS_LPS_STIM_DC | 0.081 / 0.026 / 98.01 | Chauvistré et al., *Eur. J. Immunol.* 2014; Nencioni et al., *Clin. Cancer Res.* 2007. |
| **acadesine** boosts Dendritic cell vs Dendritic cell stimulated with LPS | GSE36009_UNSTIM_VS_LPS_STIM_DC | 0.865 / 0.012 / 90.43 | Suzuki et al., *Invest. Ophthalmol. Vis. Sci.* 2012. |
| **levonorgestrel** boosts Dendritic cell vs Dendritic cell stimulated with LPS | GSE36009_UNSTIM_VS_LPS_STIM_DC | 0.103 / 0.058 / 98.47 | Quispe Calla et al., *Sci. Rep.* 2016. |
| **TAK-715** boosts Dendritic cell vs Dendritic cell stimulated with LPS | GSE36009_UNSTIM_VS_LPS_STIM_DC | 0.203 / 0.0143 / 94.72 | Miwatashi et al., *J. Med. Chem.* 2005; Xie et al., *J. Immunol.* 2003. |
| **amiodarone** inhibits Macrophage M1 vs Macrophage M2 | COATES_MACROPHAGE_M1_VS_M2 | 0.958 / 0.024 / -90.72 | Hoffman et al., *DDL2017*. 2017. |
| **piceatannol** inhibits Macrophage M1 vs Macrophage M2 | COATES_MACROPHAGE_M1_VS_M2 | 0.764 / 0.014 / -90.64 | Mahon et al., *Acta. Biomater.* 2018. |
| **AM-404** inhibits Macrophage M1 vs Macrophage M2 | COATES_MACROPHAGE_M1_VS_M2 | 0.560 / 0.165 / -91.84 | Tchantchou et al., *Neuropharmacology* 2014. |
| **doramapimod** boosts Macrophage M1 vs Macrophage M2 | COATES_MACROPHAGE_M1_VS_M2 | 0.288 / 0.15 / 92.38 | Jimenéz-Garcia et al., *Eur. J. Immunol.* 2014; Moon et al., *J. Pharmacol. Sci.*2015. |
| **staurosporine** boosts Macrophage M1 vs Macrophage M2 | COATES_MACROPHAGE_M1_VS_M2 | 0.143 / 0.718 / 93.2 | Kurian et al., *Eur. Respir. J.* 2014. |
| **tyrphostin-B44** inhibits T CD4+ stimulated with Retinoic Acid vs T CD4+ stimulated with RARA antagonist | GSE20500_RETINOIC_ACID_VS_RARA_ ANTAGONIST_TREATED_CD4_TCELL | 0.065 / 0.052 / -99.92 | Jetten. *Nature* 1980. |
| **prostaglandin-e1** inhibits mature Dendritic cell vs mature Dendritic cell stimulated with PGE2 | GSE9946_MATURE_STIMULATORY_VS_ PROSTAGLANDINE2_TREATED_MATURE_DC | 0.253 / 0.097 / -91.11 | Steinbrink et al., *Arch. Dermatol. Res.* 2000. |
| **trans-7-hydroxy-pipat** inhibits T helper 17 vs induced T regulatory | GSE14308_TH17_VS_INDUCED_TREG | 0.031 / 0.114 / -99.92 | Lu et al., *Biomed Res. Int.* 2015. |

* * * * * * * * * *
